# Proteome-wide solubility and thermal stability profiling reveals distinct regulatory roles for ATP

**DOI:** 10.1101/488163

**Authors:** Sindhuja Sridharan, Nils Kurzawa, Thilo Werner, Ina Günthner, Dominic Helm, Wolfgang Huber, Marcus Bantscheff, Mikhail M Savitski

**Author notes:** Equally contributing authors (Sindhuja Sridharan, Nils Kurzawa). Corresponding authors: Address all correspondence to or.

## Abstract

Nucleotide triphosphates (NTPs) regulate numerous biochemical processes in cells as (co-)substrates, allosteric modulators, biosynthetic precursors, and signaling molecules^1–3^. Apart from its roles as energy source fueling cellular biochemistry, adenosine triphosphate (ATP), the most abundant NTP in cells, has been reported to affect macromolecular assemblies, such as protein complexes^4,5^ and membrane-less organelles^6–8^. Moreover, both ATP and guanosine triphosphate (GTP) have recently been shown to dissolve protein aggregates^9^. However, system-wide studies to characterize NTP-interactions under conditions approximating the native cellular environment are lacking, which limits our perspective of the diverse physiological roles of NTPs. Here, we have mapped and quantified proteome-wide NTP-interactions by assessing thermal stability and solubility of proteins using mechanically disrupted cells. Our results reveal diverse biological roles of ATP depending on its concentration. We found that ATP specifically interacts with proteins that utilize it as substrate or allosteric modulator at doses lower than 500 μM, while it affects protein-protein interactions of protein complexes at mildly higher concentrations (between 1-2 mM). At high concentrations (> 2 mM), ATP modulates the solubility state of a quarter of the insoluble proteome, consisting of positively charged, intrinsically disordered, nucleic acid binding proteins, which are part of membrane-less organelles. The extent of solubilization depends on the localization of proteins to different membrane-less organelles. Furthermore, we uncover that ATP regulates protein-DNA interactions of the Barrier to autointegration factor (BANF1). Our data provides the first quantitative proteome-wide map of ATP affecting protein structure and protein complex stability and solubility, providing unique clues on its role in protein phase transitions.

---

Proteome-wide studies have improved our understanding of protein-metabolite interactions in extracts of bacteria^4^ and mammalian cells^10–13^. However, most studies utilize cellular lysates containing only the soluble proteome. To specifically assess the global roles of ATP and GTP under conditions approximating the native cellular environment, we mechanically disrupted Jurkat cells to obtain ‘crude lysates’ that retain insoluble proteins, protein condensates, and membrane proteins embedded in lipids^14–16^. In these lysates, we studied the effects of ATP and GTP on protein thermal stability on a proteome-wide scale by combining the principle of the cellular thermal shift assay^17^ with multiplexed quantitative mass spectrometry ^18^. This approach termed thermal proteome profiling (TPP) (fig. S1A) monitors how melting curves of proteins change upon treatment with a single NTP concentration^10–12,19^. Further, we used two dimensional TPP (2D-TPP) to determine the amount of NTP required to alter protein thermal stability (Fig. 1A)^13^.

**Fig. 1.**
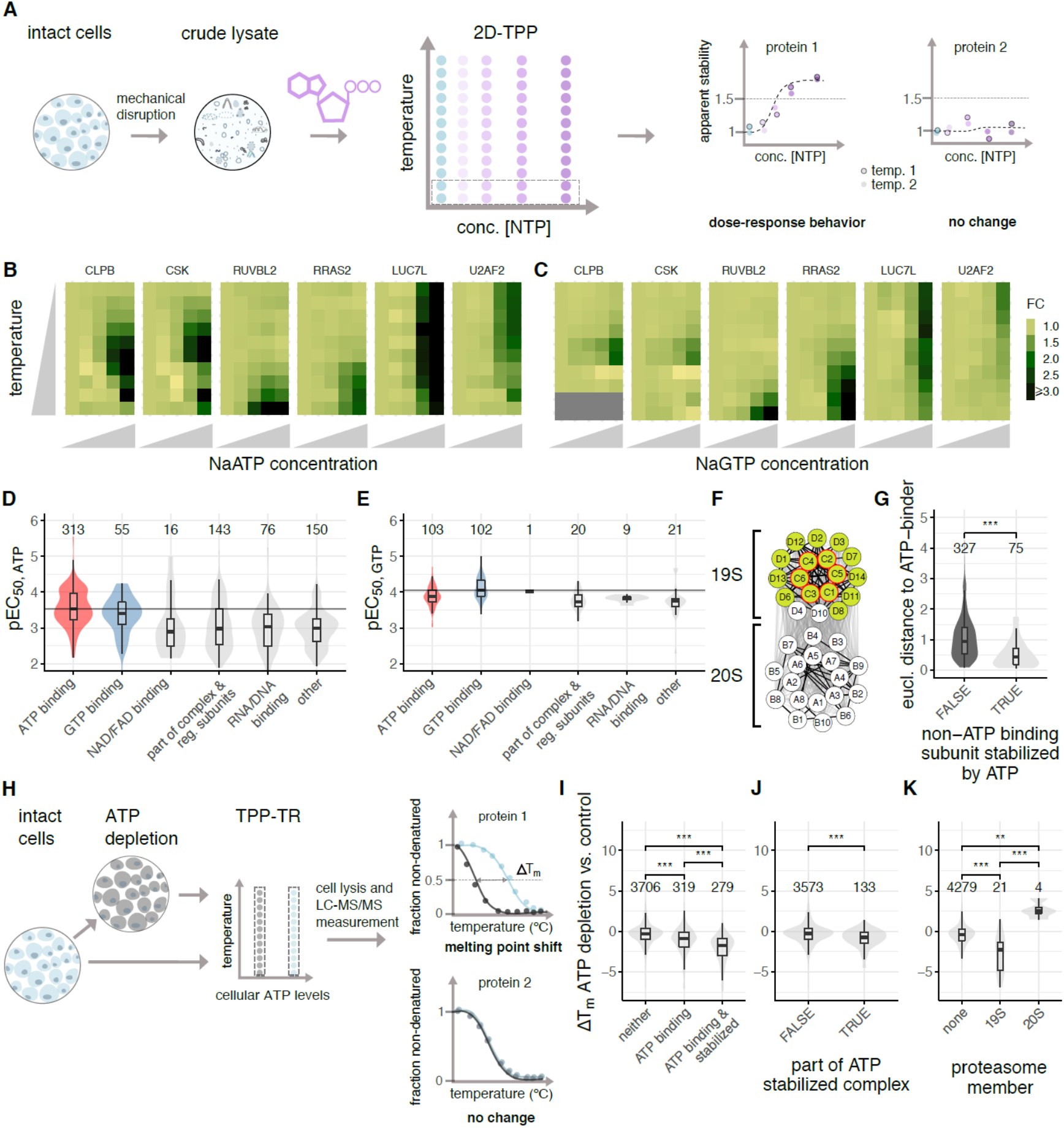
Effect of ATP and GTP on proteome thermal stability. (**A**) Experimental setup of 2D thermal proteome profiling (2D-TPP) using crude lysate system. Dotted rectangular box corresponds to one TMT10 experiment. (**B**) Heat maps showing relative fold changes (FC) of protein abundance upon treatment with ATP (0.005, 0.05, 0.5 and 2 mM) compared to untreated crude lysate (first column on each plot) with increasing temperature (y-axis: 42, 44.1, 46.2, 48.1, 50.4, 51.9, 54.0, 56.1, 58.2 and 60.1°C) (**C**) Heat maps showing relative FC of protein abundances upon treatment with GTP (0.001, 0.01, 0.1 and 0.5 mM) compared to untreated crude lysate (first column on each plot) with increasing temperature (y-axis: 42, 44.1, 46.2, 48.1, 50.4, 51.9, 54.0, 56.1, 58.2, 60.1, 62.4 and 63.9 °C) (**D-E**) Distribution of −log_10_ half-maximal effective concentration (pEC_50_) values of different classes of proteins as annotated in UniProt stabilized by ATP (D), and GTP (E). (**F**) Network diagram showing effect of ATP on proteasome stability. Nodes with green (filled) circles indicate ATP stabilized subunits, and a red outline for nodes show known ATP binding proteins of the complex. Edge thickness represents the Euclidean distance between the melting profiles of the different subunits. Thick lines are indicative of Euclidean distance less than 0.02. (**G**) Comparison of Euclidean distances between melting profiles of stabilized ATP binding complex subunits and non-stabilized non-ATP binding subunits, (left), and between stabilized ATP binding complex subunits and stabilized non-ATP binding subunits (right) within different complexes. Significance levels obtained from a Wilcoxon signed-rank test were encoded as *p < 0.05, **p < 0.01, and ***p < 0.001. (**H**) Experimental setup of TPP on cells depleted of ATP by inhibiting glycolysis and oxidative phosphorylation with a combination of 10 mM 2-deoxyglucose and 1 nM Antimycin-A. Dotted rectangular box corresponds to one TMT10 experiment. (**I**) Changes in melting points of non-ATP binding proteins, ATP binding proteins, and ATP binding proteins that were stabilized by ATP in the crude lysate experiment upon ATP depletion in cells. Significance levels obtained from a Wilcoxon signed-rank test were encoded as *p < 0.05, **p < 0.01, and ***p < 0.001. (**J**) Changes in melting points of protein complex subunits stabilized by ATP in the crude lysate experiment (right), and all other proteins (left) upon ATP depletion in cells. Significance levels obtained from a Wilcoxon signed-rank test were encoded as *p < 0.05, **p < 0.01, and ***p < 0.001. (**K**) Changes in melting points of 19S and 20S proteasome subunits, and all other proteins upon ATP depletion in cells. Significance levels obtained from a Wilcoxon signed-rank test were encoded as *p < 0.05, **p < 0.01, and ***p < 0.001.

Comparison of TPP experiments performed at 2 mM ATP or vehicle revealed similar melting point shifts in crude and gel-filtered lysates (fig. S1B, table S1) confirming that crude lysate is a suitable system for detecting protein-metabolite interactions. Intracellular ATP concentrations are highly variable in cells (1-10 mM)^20^, and GTP levels are approximately 5-fold lower than ATP^21^. We found that melting point shifts induced by 2 mM ATP and 0.5 mM GTP in crude lysate were selective for known ATP and GTP binding proteins, respectively (fig. S1C, table S2) and were used as maximum concentrations in dose-dependent studies by 2D-TPP (Fig. 1 A-C). The 2D-TPP analysis uncovered 753 proteins with increased thermal stability in the presence of ATP, out of the 7549 identified proteins at a 1% false discovery rate (FDR) (fig. S2, see methods). We determined the affinity of NTP to a protein, as the half-maximal effective concentrations of protein stabilization (EC_50_, pEC_50_ = −log_10_(EC_50_)). The largest and most potently affected group of proteins that were stabilized by ATP represented annotated ATP-binding proteins (315 proteins), validating our approach. A small subset of GTP-binding proteins also showed increased thermal stability with added ATP (55 proteins) (Fig. 1D and table S3). Moreover, ATP reduced the thermal stability of 151 proteins, which were enriched for RNA binding proteins, including ribosomal proteins (p-value < 0.001), in line with previous observations in *E. coli* ^4^.

The 2D-TPP experiment with GTP in crude lysate revealed increased thermal stability for 256 out of 7618 identified proteins, with the two largest groups being GTP-(102 proteins) and ATP-(103 proteins) binding proteins (Fig. 1E, table S3), whereas 9 proteins showed lower thermal stability. Proteins containing phosphate binding-loops (P-loops) were stabilized by ATP or GTP, as expected^22^. We found that the residues flanking the Walker-A motif have a substantially more constrained composition in GTP-stabilized proteins compared to ATP-stabilized ones (fig. S3 A-D). Proteins that were stabilized by both NTPs included a small subset of kinases and proteins with P-loops (AAA-type ATPases and small–GTPases) (fig. S3E-H, table S3 and S4). Amongst these were several proteins known to interact with both metabolites, such as Uridine-cytidine kinase (UCK1 and UCK2), casein kinase (CSNK1A1, CSNK1D, CSNK2A1) and succinyl CoA synthetase (SUCLA2 and SUCLG2)^23–27^. Importantly, we identified proteins not previously reported to have this property, such as, ATP binding cassette super family proteins (ABCE1, ABCF1, etc.), Caseinolytic peptidase B protein homolog (CLPB), Tyrosine-protein kinase (CSK), RuvB-like ATPase (RUVBL2), Rab-GTPases (RAB21, RAB10, RAB2A), Ras-related protein R-Ras2 (RRAS2), and phosphofructokinase (PFKL) (Fig. 1B and C, fig. S3I-J, table S3). In summary, we observed crosstalk between ATP- and GTP-induced thermal stabilization, confirming previously known ATP/GTP-binding proteins and uncovering potential new ones, and showed that ATP- and GTP-binding preferences reflect different sequence constraints on the Walker-A motif flanking residues.

Similar to a recent system-wide study in *E. coli*^4^, more than half of the ATP-affected proteins were not annotated in UniProt as ATP or GTP binders. These proteins required higher ATP concentrations for stabilization and were enriched for NAD/FAD, DNA, and RNA binding proteins, as well as for components of annotated complexes and regulatory subunits (Fig. 1D). We observed thermal stabilization of several NADH dehydrogenases, such as GAPDH, IDH1, IDH2, MDH1, MDH2 and PDHX, which have been reported to be inhibited by ATP^28–31^. This may suggest that higher ATP levels in cells could cause feedback inhibition of these enzymes. Looking specifically at complexes, we observed preferential stabilization of non-ATP-binding subunits that form complexes with at least one ATP-binding subunit (p-value < 0.001). This suggests that thermal stabilization of an ATP-binding complex subunit can propagate to proximal non-ATP-binding subunits. The proteasome provides a case-in-point: the ATP-binding PSMC subunits and the majority of the non-ATP-binding PSMD subunits of the 19S regulatory particle were stabilized by ATP. In contrast, the thermal stability of the 20S core particle, which is devoid of ATP-binding subunits, was unaltered. Comparing the melting curves of proteins by calculating the average Euclidean distance^32^, we observed that both 19S and 20S subunits exhibit significant co-melting behaviors (p < 0.001) that were different between the particles, indicating that the ATP induced stabilization propagates only to subunits that are in physical proximity (Fig. 1F). Global analysis revealed that similar melting behaviors of the stabilized non-ATP binding subunits and the stabilized ATP binding subunits within protein complexes is a general characteristic of ATP-induced complex stabilization (Fig. 1G and S4).

To investigate whether our observations made in crude lysates are relevant in intact cells, we reduced ATP levels in Jurkat cells by inhibiting glycolysis and oxidative phosphorylation, using 2-deoxyglucose and Antimycin-A, respectively (fig. S5A-B)^33^ and compared the thermal stability profiles of untreated and ATP-depleted cells (Fig. 1H). Indeed, among the 5199 proteins identified, the ATP-binding proteins (279) that were stabilized by addition of ATP to crude lysates showed the most pronounced thermal destabilization in ATP-depleted cells (Fig. 1I, table S4). Likewise, protein complex subunits that were stabilized by ATP in lysate were destabilized in ATP-depleted cells (Fig. 1J and K, table S4). Finally, the proteins that were destabilized by ATP in lysates showed significant thermal stabilization upon cellular ATP depletion (p-value < 0.001) (fig. S5C, table S4). Thus, we conclude that our findings with crude lysates reflect physiologically relevant roles of ATP.

Crude lysate 2D-TPP experiments revealed several proteins that apparently increase in abundance with higher ATP and GTP concentrations, already at 42°C (e.g., LUC7L, Fig. 1 B and C). This is indicative of increased solubility rather than thermal stability^13^, and supports the recently discovered function of ATP as a biological hydrotrope, which solubilizes hydrophobic molecules in aqueous solution^9^. To systematically assess the role of ATP in proteome solubility, we devised an experimental strategy that we termed Solubility Proteome Profiling (SPP), which uses multiplexed mass spectrometric analysis to quantify the concentration-dependent effect of a molecule (e.g., ATP) on the soluble and insoluble populations of individual proteins, on a proteome-wide scale (Fig. 2A). Briefly, crude cell lysate was divided into 10 aliquots, of which nine were treated with either vehicle or increasing concentrations of ATP (0.1 to 10 mM), followed by solubilization of membrane-or organelle-bound proteins with the mild detergent NP40. A tenth, vehicle-treated aliquot was solubilized with the strong detergent SDS (1%). The ratios between the SDS and NP40 solubilized samples determined for each protein inform on the fractions that exists in an insoluble vs. a soluble state, where a high ratio suggests a higher insoluble fraction ^34^. By determining the half-maximal solubilization concentrations (pEC_50_s), our experiments reveal the differential susceptibility of proteins to solubilization by ATP.

**Fig. 2.**
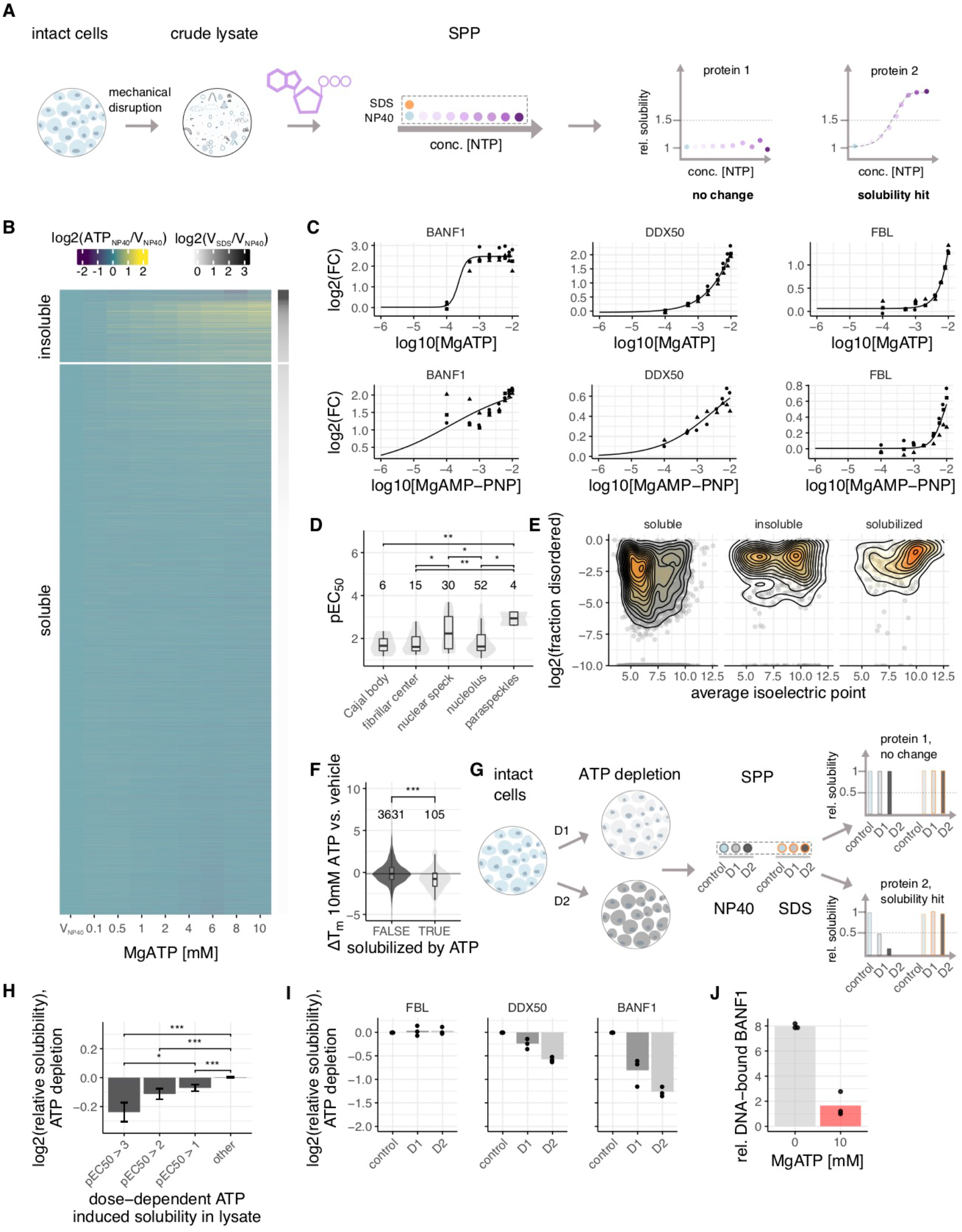
Solubility proteome profiling characterizes broad solubilizing effects of ATP on the proteome. (**A**) Experimental setup of solubility proteome profiling (SPP). Cells were lysed by mechanical disruption and the resulting crude lysate was divided into ten aliquots. Two crude lysate aliquots were treated with vehicle and eight aliquots with different concentrations of a molecule (e.g. ATP). One vehicle treated aliquot and eight molecule treated samples were solubilized with a mild detergent (NP40) and the other vehicle control was solubilized with strong detergent (SDS) All samples were digested with trypsin, labeled with different TMT10 isotope tags and analyzed by LC-MS/MS, dotted rectangular box corresponds to one TMT10 experiment. Proteins that showed at least 50% increase in abundance in NP40 processed molecule treatment (at least in one concentration) compared to NP40 processed vehicle, were fitted with sigmoidal dose-response curves and pEC_50_ values were calculated. (**B**) Heat map representation of proteome solubility. The grey scale represents the log_2_ ratio between NP40 and SDS processed vehicle condition. The color scale represents the log_2_ ratio between NP40 processed ATP treated and vehicle treated conditions. (**C**) Solubility profiles of BANF1, DDX50, and FBL, following ATP (upper panel) and non-hydrolyzable ATP (AMP-PNP) (lower panel) treatment, y-axis represents the log_2_ ratio of NP40 processed ATP or AMP-PNP treated and vehicle treated conditions. (**D**) Distribution of −log_10_ half-maximal effective concentration (pEC_50_) values of ATP-solubilized proteins allocated to different membrane-less organelles. Significance levels obtained from a Wilcoxon signed-rank test were encoded as *p < 0.05, **p < 0.01, and ***p < 0.001. (**E**) 2D density contours of log_2_ fraction disorder vs. average isoelectric points of soluble and insoluble proteins, and of those solubilized by ATP. (**F**) Melting point differences of proteins solubilized by ATP (from SPP data) compared to all other proteins measured by TPP between 10 mM ATP and vehicle treated crude. Significance levels obtained from a Wilcoxon signed-rank test were encoded as *p < 0.05, **p < 0.01, and ***p < 0.001. (**G**) Experimental setup of SPP in ATP depleted cells. Cells were depleted of ATP by inhibiting glycolysis and oxidative phosphorylation with two combinations of 2-deoxyglucose (2DG) and Antimycin-A (AA) (D1: 0.1 nM AA and 1 mM 2DG, D2: 1 nM AA and 10 mM 2DG). The untreated cells and the cells from the two ATP depleted conditions were divided into two aliquots each, of which one was solubilized with NP40 while the other using 1% SDS. All samples were digested with trypsin, labeled with different TMT10 isotope tags and analyzed by LC-MS/MS. (**H**) Change in solubility calculated for proteins binned according to their propensity for solubilization with ATP in crude lysate (pEC_50_ x-axis), following ATP depletion in cells. Significance levels obtained from a Wilcoxon signed-rank test were encoded as *p < 0.05, **p < 0.01, and ***p < 0.001. (**I**) Change in solubility of FBL, DDX50, and BANF1 following ATP depletion in cells, measured by calculating the log_2_ ratios of NP40 processed untreated and ATP depleted conditions. (**J**) DNA binding propensity of BANF1 in the presence of ATP. Recombinantly expressed and purified BANF1 was incubated with biotinylated double stranded DNA in the presence and absence of ATP. The DNA-bound fraction of BANF1 was pulled down using streptavidin beads and measured using quantitative mass spectrometry. Rel. DNA bound fraction of BANF1 was calculated as the ratio of intensity of protein in the presence and absence of DNA to correct for non-specific binding of protein to beads.

To investigate whether ATP has a prominent role as a biological hydrotrope, we examined the insoluble proteome. Proteins that were at least 50% more abundant in the SDS-processed vehicle condition compared to the NP40-processed vehicle condition were defined as part of the insoluble proteome^34^. It should be noted that most of these proteins have a soluble subpopulation (Fig. 2B, table S5). We observed ATP dependent solubilization for 188 proteins out of the 760 proteins that were identified as insoluble in our experiments, and calculated pEC_50_s for the dose dependent solubilization of these proteins (Fig. 2B, table S5). These data showed that ATP solubilized almost 25% of the insoluble proteome. Intriguingly, only 23% of the solubilized proteins were annotated as ATP binders whilst the majority (54%) were annotated as part of membrane-less organelles (fig. S6A, table S5). Further, we found that proteins exhibited differential susceptibility for solubilization. For example, proteins such as myosin (MYO1G) which is known to dissociate from actin in the presence of ATP^35^, BANF1 and DDX50 were potently solubilized by ATP with sub-micromolar pEC_50_s, whereas other proteins, such as FBL, NOP56 and EIF3H were only solubilized at much higher ATP concentrations, (Fig. 2C, fig. S6B, and table S5). Strikingly, the susceptibility of proteins for being solubilized by ATP depended substantially on their localization in different membrane-less organelles. For example, proteins annotated as part of nuclear speckles solubilized, on average, at lower ATP concentrations than proteins annotated as part of the nucleolus (Fig. 2D). In general, the ATP-solubilized proteins were significantly enriched in disordered regions and had significantly higher isoelectric points than the rest of the proteome (Fig. 2E, fig. S6 C and D, and table S5). The solubilizing properties of ATP were mimicked by GTP (fig. S7, table S6), as well as by the non-hydrolysable analog of ATP (AMP-PNP), (Fig. 2C, fig. S8, and table S7) showing on a proteome-wide scale that ATP hydrolysis is not the main factor for ATP-mediated solubilization and suggesting that previous findings based on fluorescently tagged nucleolus markers FBL and NPM1 cannot be generalized^36^.

Furthermore, we used TPP to investigate how the addition of 10 mM ATP affected the thermal stability of solubilized proteins (table S8) and observed a significant trend for thermal destabilization compared to the vehicle condition (Fig. 2F). This suggests that the ATP solubilized fractions of these proteins tended to have lower thermal stability than the soluble ones which may be due to distinct interactions or post-translational modifications in the two populations.

To extend our analysis of the role of ATP on proteome solubility to cells, we again depleted ATP from Jurkat cells with 2-deoxyglucose and Antimycin-A (Fig. 2G). We identified 107 proteins with decreased solubility—14% of the insoluble proteome—in the ATP-depleted cells compared to vehicle condition (table S9, fig. S9A). Since ATP depletion is expected to affect several homeostatic processes, our analysis primarily sought to assess the changes in solubility in cells of proteins that gained solubility in the crude lysate system. (Fig. 2G, fig. S9B and C, table S9). In general, proteins solubilized at lower ATP concentrations in crude lysate had a stronger decrease in solubility upon cellular ATP depletion (Fig. 2H). For example, FBL, which is solubilized at higher ATP concentrations in crude lysate, showed no decrease in solubility in cells upon ATP depletion (Fig. 2I). In contrast, DDX50 and BANF1, two proteins that were solubilized with low concentrations of ATP in crude lysate, displayed decreased solubility following ATP depletion in cells (Fig. 2I). We thus conclude that the solubilizing effects of ATP are also relevant in cellular systems.

Following the global analysis, we set out to understand the solubility changes observed in one of the highly ATP-solubilized protein, Barrier to autointegration factor (BANF1)—which is not annotated to be part of any membrane-less organelles. BANF1 is a non-specific DNA binding protein which transiently crossbridges anaphase chromosomes to promote assembly of a single nucleus^37^. We hypothesized that ATP may affect DNA binding abilities of BANF1 and hence tested if purified BANF (Fig. S10) bound double stranded DNA in the presence and absence of ATP. We observed that the ability of BANF1 to interact with DNA is nearly six times lower in the presence of 10 mM ATP (Fig. 2J). This suggests that ATP solubilizes BANF1 by preventing its binding to DNA, and thus represents an example of ATP regulating protein-DNA interactions. ATP depletion is known to cause chromatin compaction ^38^. Our data shows that BANF1 loses solubility upon ATP depletion, suggesting its interaction with chromatin. This suggests a role of BANF1 in compacting DNA at low ATP levels.

In crude lysates a few proteins became less soluble with increasing ATP concentration, with two prominent examples being IMPDH1 and NUCKS1, (Fig. 3A, table S5). In both cases, ATP hydrolysis is likely to play a role, since neither protein decreased in solubility following addition of the non-hydrolyzable ATP analog AMP-PNP (fig. S8E). ATP was reported to act as an allosteric modulator of recombinant IMPDH1, driving the formation of a filamentous structure^39^, in agreement with our observations on endogenous IMPDH1. NUCKS1 is predicted to be a highly disordered (99%) chromatin associated protein, which is necessary for DNA repair by homologous recombination^40^. Single nucleotide polymorphisms of NUCKS1 have been linked through genome wide association studies to Parkinson’s, a protein aggregation-related disease^41^. The melting behavior of NUCKS1 in TPP experiments treated with 10 mM ATP followed a striking pattern, with a sharp increase of soluble protein abundance peaking at approximately 58°C. At higher temperatures a temperature-dependent decrease of soluble protein was observed, with the melting curve matching that determined in the vehicle condition (Fig. 3B). This may suggest that the ATP related metabolism induces NUCKS 1 to transition into two-phases, which is observed as loss in solubility of the protein. Upon heating, the NUCKS1 assembly transitions into a single phase at 58°C, indicative of its critical transition temperature^42^. In contrast, IMPDH1 exhibits a more gradual increase in solubility at lower temperatures, and shows a substantially higher thermal stability, indicating that the ATP binding^39^ leads to a more stable conformation (Fig. 3B). Finally, we asked ourselves if proteins that have insoluble subpopulations under physiological conditions (without addition of ATP) would in general show some increase in solubility upon heating. In order to answer that question, we looked specifically at maximal log_2_ fold changes across the heating temperatures—calculated using 37°C as reference—observed for melting curves of proteins that have an insoluble subpopulation in the crude lysate (ratio of SDS/NP40 > 1.5). Indeed, for this group of proteins we saw a significant increase in the amount of protein that became soluble upon heating (Fig. 3C), indicating temperature-dependent phase transitions on a proteome-level. These observations suggest that the combination of thermal stability and solubility could potentially be used to characterize phase transitions on a system-wide scale.

**Fig. 3.**
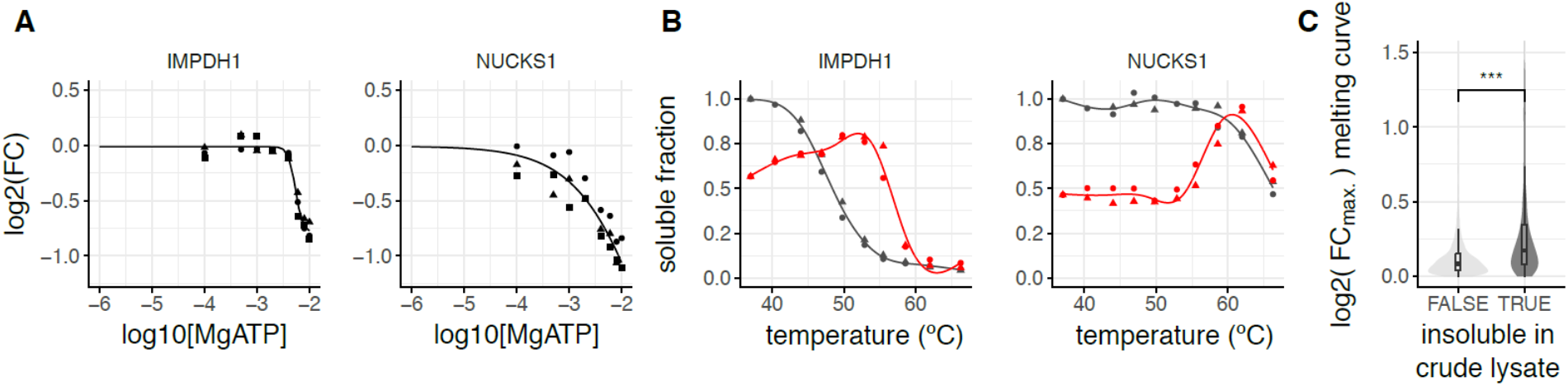
Solubility proteome profiling identifies proteins decreasing in solubility upon ATP addition. (**A**) Solubility profile for IMPDH1 and NUCKS1, fold change (FC) in y-axis represents the ratio of NP40 processed ATP treated and vehicle treated conditions. (**B**) Melting curves for IMPDH1 and NUCKS1 from TPP experiments in crude lysates treated with vehicle (black) or 10mM ATP (red) and corrected for solubility changes at 10 mM ATP using SPP data. (**C**) Maximal log_2_ fold changes observed for melting curves of insoluble versus soluble proteins in untreated crude lysate. Significance levels obtained from a Wilcoxon signed-rank test were encoded as *p < 0.05, **p < 0.01, and ***p < 0.001.

Together, our TPP, 2D-TPP and SPP data revealed concentration-dependent roles of ATP. At low concentrations (below 500 μM), ATP mainly acted as a substrate for nucleotide-binding proteins. At intermediate concentrations (between 1 – 2 mM), broad effects of ATP on the stability of protein complexes were observed. At higher concentrations (greater than 2mM) ATP has solubilizing effects, primarily on disordered proteins with positive charge that bind nucleic acids such as DNA and RNA (Fig. 4). Our SPP analysis revealed that ATP serves as a hydrotrope to a substantial fraction of the insoluble proteome that is enriched for components of membrane-less organelles. These data strongly suggest that ATP regulates phase separation not only by regulating protein activities^6,8^, but also by directly promoting protein solubility. Taken together, the ATP-dependency exhibited on a proteome-level suggests that metabolic fluctuations, such as nutrient deprivation, that affect ATP concentrations in cells, may be sensed as the altered solubility status of nucleic acid binding proteins. Our data provides a unique map of the subproteomes affected in their solubility and thermal stability by ATP and will facilitate the discovery of new ATP-binding proteins, components of membrane-less organelles and regulators of phase separation. The SPP approach described herein can be applied in future studies to assess the effects of other molecules and cellular perturbations on protein solubility in different disease-relevant cell types, to study regulation of aggregation-susceptible proteins and organelles.

**Fig. 4.**
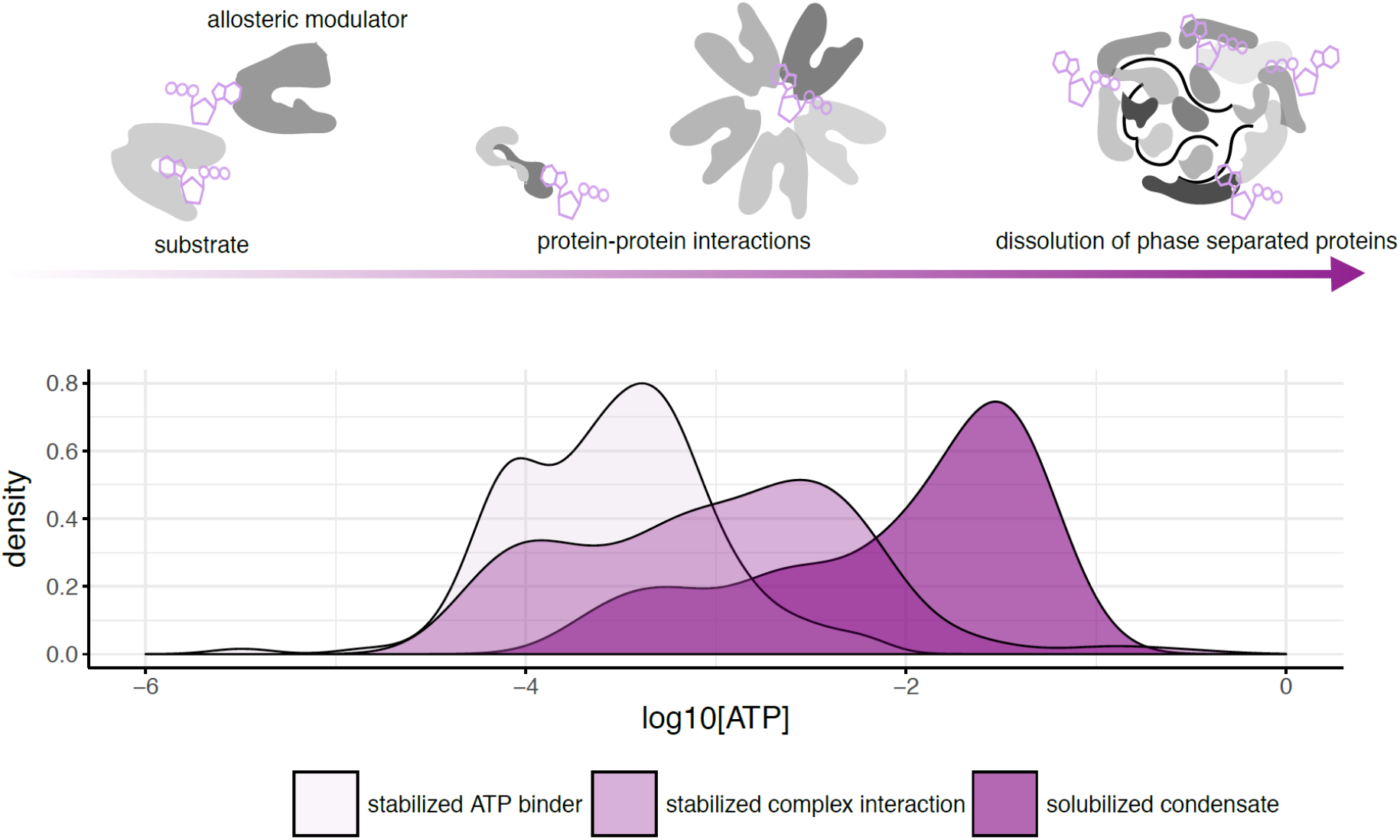
Concentration dependent proteome-wide effect of ATP. Density plots of pEC_50_s of ATP-stabilized ATP-binding proteins, complexes, as well as of proteins solubilized by ATP as measured by 2D-TPP and SPP technologies. Despite a clear difference in the distributions, there is a substantial overlap, showing that changes in ATP concentrations will have simultaneous effects on the stability of ATP-binding proteins and complexes as well as on the solubility of disordered proteins.

## Author Contributions

S.S., N.K., M.B., and M.M.S. conceptualized the study, S.S., T.W., M.B., and M.M.S. designed the experiments, S.S., T.W., I.G., and D.H. performed the experiments N.K., W.H., M.B., and M.M.S. designed and implemented data analysis strategies, S.S., N.K., M.B. and M.M.S. analyzed the data, S.S., N.K., M.B. and M.M.S. wrote the manuscript.

## Acknowledgments

We thank Andre Mateus and Life Science Editors for insightful discussions and proofreading of the manuscript, Natalie Romanov for supplying and helping with protein complex and disorder annotations; the Proteomics Core Facility at the European Molecular Biology Laboratory (EMBL) for expert help; K. Remans, J. Scheurich and J. Flock from Protein Expression and Purification Core Facility at EMBL for help in recombinant expression and purification of BANF1, Friedrich Reinhard and Christina Rau at Cellzome-GSK for help with the crude lysate protocol and J. Stuhlfauth and N. Garcia-Altrieth at Cellzome-GSK for cell culture; M. Jundt, K. Kammerer, M. Klös-Hudak, M. Paulmann, and T. Rudi at Cellzome-GSK for expert technical assistance.

